# Organoid-evaluable clinical biomarkers predict drug responses and guide new breast cancer therapies

**DOI:** 10.1101/2025.08.24.671810

**Authors:** Tam Binh V. Bui, Denise M. Wolf, Michael C. Bruck, Kaitlin Moore, Jessica Lien, Sarah D.W. Choi, Shruti Warhadpande, Amirabbas Parizadeh, Deborah Dillon, Beth Overmoyer, Filipa Lynce, I-SPY 2 Investigators, Isaac J. Nijman, Boudewijn M.T. Burgering, Isaac S. Harris, Laura J. Esserman, Laura J. van ‘t Veer, Jennifer M. Rosenbluth

**Author notes:** **Corresponding author** Jennifer M. Rosenbluth, MD, PhD Department of Medicine, University of California, San Francisco. **Declaration of interests** Laura J. van ‘t Veer is a part-time employee and stockholder of Agendia. Laura J. Esserman reported that she is an uncompensated board member of the Quantum Leap Healthcare Collaborative, which sponsors the I-SPY trial. No potential conflicts of interest were disclosed by the other authors.

## Abstract

Poor therapeutic response in subsets of breast cancer (BC) patients poses an ongoing challenge. Here, we present a biomarker-guided characterization of 44 patient-derived BC organoids, with the aim of modeling resistant disease with greater fidelity and developing an in-vitro system grounded in clinical data for testing alternative treatment strategies. We utilized patient transcriptomic and outcome data from the I-SPY2 clinical trial to develop predictive models of response to a range of therapies, using only organoid-detectable biomarkers as input. A model predicting response to veliparib-platinum chemotherapy (VP) in triple-negative BC (TNBC) was validated in organoids, showing that in vitro drug responses matched predictions from the patient data-derived model. A drug screen in VP-resistant TNBC organoids identified combination treatments that overcame resistance to cisplatin, including pro-apoptotic therapies. This demonstrates that gene expression-based resistance models derived from patient data can be successfully modeled in organoids that can then be used for therapeutic evaluation.

## Introduction

An important challenge in cancer research is developing robust and representative research models that can accurately recapitulate the complexity of the disease(1). While traditional cell lines have provided valuable insights, they often fail to capture the full spectrum of cancer subtypes and fail to maintain the wide range of genetic, phenotypic, and molecular heterogeneity seen in the original tumors (2), in part because the rate of successful establishment of cell lines from primary tumor specimens remains low. Patient-derived organoids, in contrast, can be established with higher efficiency and can recapitulate important histological and genetic features of the tumor tissue (3,4). Another major area of recent advancement is the development of clinical biomarkers that can predict patient treatment response and support clinical decision-making (5). It remains unknown, however, whether and to what extent clinical gene expression-based biomarkers and predictive models derived from and predictive in patients can be applied to organoid models. While key studies have evaluated individual or small numbers of markers in cancer organoids(6,7), it is largely undetermined which gene expression-based biomarkers can be modeled in organoids.

This is particularly relevant in breast cancer, where drug development and clinical trial progress in recent years have seen an increased number of treatment options available to patients, creating a need for better biomarkers and subtyping schemas to identify and predict the populations who will benefit. The ongoing, multi-center phase 2 neoadjuvant chemo-/targeted-therapy I-SPY2 platform trial was designed to rapidly screen promising experimental agents and identify improved treatment regimens in women with early-stage, high molecular risk breast cancer, as well as to identify and propose potential biomarkers of response and resistance. The primary endpoint of the trial is pathologic complete response (pCR), defined as the absence of invasive breast cancer in the breast and regional nodes at the time of surgery due to the complete eradication of the cancer cells by the preceding therapy. The trial developed the Response Predictive Subtypes (RPS) (4), a breast cancer classification schema designed to better segregate tumors based on anticipated responses to therapies in a modern treatment landscape, including immunotherapy, PARP-inhibition, platinum drugs, and dual-HER2 inhibition, by combining Immune, DNA repair deficiency (DRD), and Luminal-like biological phenotypes with hormone receptor (HR) and HER2 status (5). Importantly, the efforts to improve breast cancer classification have identified subsets of patients who do not respond to neoadjuvant targeted and/or chemotherapy. Therefore, a deeper understanding of the molecular mechanisms driving breast cancer resistance is needed to inform the development of more effective therapeutic strategies that are targeted to patients who do not respond to existing treatments. Organoids hold potential for identifying new personalized treatment strategies for cancer patients, particularly if they can be shown to stratify patients using the same biomarkers that can stratify patients in the clinic and in clinical trials.

In this paper, we evaluated a biobank of early-stage invasive breast cancer organoids characterized using I-SPY2 predictive biomarkers as a tool to model and study mechanisms of therapy resistance. As a proof-of-concept, we developed organoid-tailored predictive models derived from patient biomarker and clinical data in the I-SPY2 clinical trial and then used these to predict treatment sensitivity and resistance in subtype-matched organoid cultures. We validated resistant organoid cultures that were then selected and used for compound screening to identify new treatment options for biomarker-identified resistant tumors. Previously, large-scale drug screening in patient-derived colon cancer and glioblastoma organoids, and PDX-derived breast cancer organoids has been reported to be feasible and useful (8–19), though this study is the first to report a high-throughput drug screen in breast cancer organoids that were derived directly from patient specimens.

## Results

### Organoids recapitulate most breast tumor subtypes

We hypothesized that organoids retain the expression of clinically relevant biomarkers and gene signatures representing key biological pathways in cancer, and that these biomarkers can be used to identify treatment-resistant organoids, which can be further used for drug testing to overcome resistance (**Figure 1A**). To this end, we evaluated bulk RNA-Sequencing data from the Hubrecht Organoid Technology (HUB) Living BC Organoid Biobank (3), which contained 26 early-stage breast tumor organoids across all receptor subtypes of breast cancer (HR+HER2-n=17, triple negative (TN) n=5, HR+HER2+ n=2, HR-HER2+ n=2; Hormone Receptor (HR) meaning either Estrogen Receptor (ER) or Progesterone Receptor (PR) positive) (**Table 1**). To enable interpretation of the HUB organoid data in a larger (clinical) breast cancer context, we quantile-normalized the organoid gene expression data to bulk RNA sequencing data from TCGA breast tumor tissue samples, resulting in an integrated dataset for further analysis.

**Figure 1.**
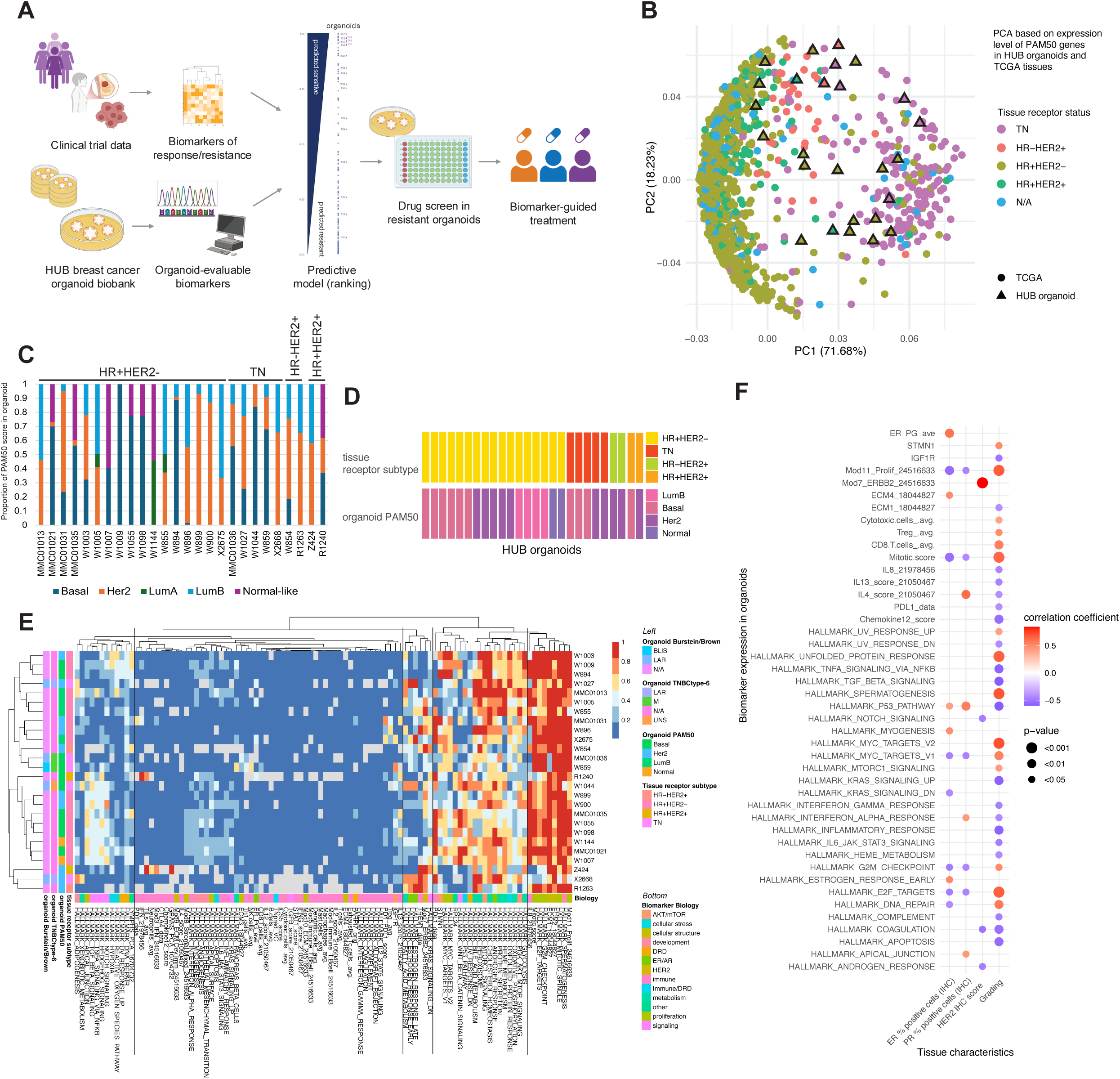
Breast cancer subtypes and clinical biomarkers can be modeled in breast cancer organoids. A. Graphical overview of the approach to model clinical trial data in organoids and identify new biomarker-guided combination therapies. HUB = Hubrecht Organoid Technology. B. Principal component analysis (PCA) showing that organoids (triangles) cluster with TCGA tissue samples (circles) based on expression levels of PAM50 genes. Colors indicate breast cancer subtypes. Axes include the percentage of variance explained by each principal component (PC). TCGA = The Cancer Genome Atlas. TN = triple negative. N/A = not available. C. Distribution of PAM50 scores components for each organoid in the HUB biobank, showing heterogeneity in each case, with the highest score resulting in the final subtype call. TN = triple negative. LumA = luminal A, LumB = luminal B. D. Subtyping of HUB organoids based on gene expression is shown for PAM50 compared to the receptor subtype of the original tumor specimen. TN = triple negative, LumB = luminal B. E. Heatmap showing the relative expression of gene expression biomarkers from the Molecular Signatures Database (MSigDB) HALLMARK set and 50 key biomarkers from the I-SPY2 clinical trial in HUB organoids. Breast cancer organoids show variable expression of biomarkers reflecting key biological features of tumors (clustered on the right, indicated by top color bar). Most but not all immune cell-based signatures (clustered on center-left) are not expressed in organoids. BLIS = basal-like immunosuppressed, LAR = luminal androgen receptor, M = mesenchymal, LumB = luminal B, TN = triple negative, UNS = a subtype could not be selected by the algorithm, N/A = not applicable (sample is not triple-negative), DRD = DNA repair deficiency, ER/AR = estrogen receptor/androgen receptor. F. Dot plot showing results of Pearson correlation analysis between expression scores of selected biomarkers in organoids (y-axis) and clinical features of the corresponding tumors (x-axis). ER = estrogen receptor, PR = progesterone receptor, IHC = immunohistochemistry.

**Table 1.**
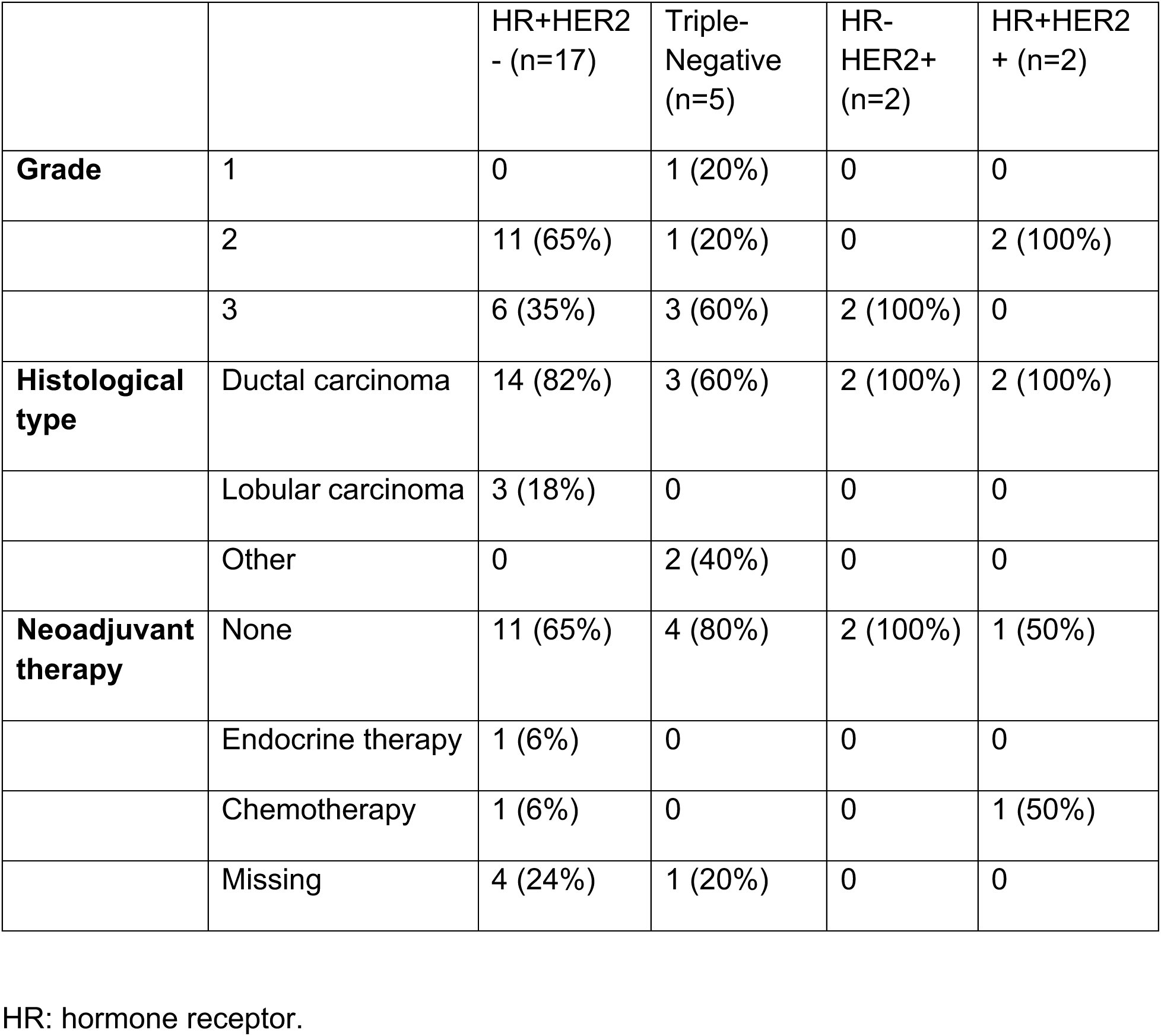
Summary of clinical characteristics of HUB organoids included in the gene expression analysis (n=26).

Principal component analysis based on the expression of PAM50 genes revealed that organoids largely cluster with TCGA samples of the same receptor subtype (**Figure 1B**), with one exception: HR+HER2-organoids are distributed more loosely across the TCGA samples, possibly due to the loss of ER signaling in organoid culture, as HUB organoids were cultured using a version of organoid medium not supplemented with estrogen. This is also reflected by the distribution of PAM50 scores in the HR+HER2-organoids (**Figure 1C-D**), with some distributions seeming to represent decreased luminal phenotype, and other distributions potentially reflecting intra-tumoral heterogeneity. Generally, our subtype predictions matched well with the previously published calls for this data set (>85% close match), with the possible exception of luminal A-classified organoids (3). Loss of ER signaling with extended culturing is an ongoing point of attention in the field, with multiple recent efforts to restore ER signaling in organoid culture (20–25).

### Patient-derived organoids retain expression of most cancer-related gene signatures

We then analyzed the expression of the 50 Molecular Signatures Database (MSigDB) Hallmark gene sets (26) by GSVA and a selection of 50 gene signatures representing key cancer signaling pathways from the I-SPY2 clinical trial, many of which correlate with treatment sensitivity and resistance in patients. We analyzed relative expression scores in HUB organoids representing the pth quantile compared to receptor-subtype matched organoids and TCGA tissues (scored between 0 and 1). Organoids in the biobank showed variable expression of ∼75% of the tested biomarkers, recapitulating key biological features of tumors, including DNA-damage response, Akt/mTOR signaling, and Myc/proliferation signatures (**Figure 1E**). While most immune signatures were negative in organoids, likely explained by the loss of immune cells after passaging of organoids (e.g., B cells and Hallmark_IL2_STAT5_signaling), several other immune signatures were detectable in organoids, likely reflecting epithelial expression of those predictive markers (e.g., PD1 and IL13).

### Organoid gene expression is consistent with clinical/pathologic tumor features

We next sought to determine the extent to which gene expression signatures measured in an organoid culture correlate with the clinical features of the patient’s tumor from which it was derived. The expression of estrogen receptor signaling-related biomarkers (ER_PG_ave, HALLMARK_ESTROGEN_RESPONSE_EARLY) in organoids was significantly correlated with ER expression levels in the tumor (% positive cells by immunohistochemistry (IHC), correlation coefficients: 0.595 and 0.511, respectively, p-value <0.05), whereas the HER2 biomarker (Mod7_ERBB2) in organoids was strongly correlated with HER2 tumor IHC-score (correlation coefficient 0.842, p-value <0.001) (**Figure 1F**). Proliferation scores (Mod11_Prolif, Mitotic score, MYC, G2M Checkpoint, E2F) were negatively correlated with ER staining and PR staining (p<0.05), but strongly positively correlated with tumor grade, as expected. Surprisingly, high expression of immune and apoptosis signatures (including HALLMARK_IL6_JAK_STAT3_SIGNALING, HALLMARK_INFLAMMATORY_RESPONSE, and HALLMARK_APOPTOSIS) was negatively correlated with tumor grade (p<0.05).

### Generation of a predictive model for organoid treatment response using biomarkers and patient response data from the I-SPY2 clinical trial

After confirming that clinically relevant biomarkers were variably expressed across the organoids and correlated with clinical features of tumors, we hypothesized that these tissue biomarkers could be used to match organoids to drug sensitivity and resistance profiles. To this end, we first sought to generate a predictive model using patient data that we could test using the organoids.

To build this model, we assessed qualifying I-SPY2 biomarkers associated with pathologic complete response (pCR) in the I-SPY2-990 clinical trial population (**Figure 2A-B**). The predictive values of biomarkers to predict pCR was evaluated in 987 patient samples across the first ten treatment arms of the trial (one control arm with standard chemotherapy and nine experimental arms with agents targeting a wide variety of pathways, including PARP inhibitors, AKT inhibitors, DNA-damaging agents, angiogenesis inhibitors, immunotherapy, small-molecule pan-HER2 inhibitors, and dual-HER2-targeting agents) and across all receptor subtypes (the “I-SPY990” dataset). We first filtered the MSigDB Hallmark/I-SPY signature set of 100 signatures to include only the 23 signatures that were previously validated in CLIA-certified labs and associated with clinical outcome (termed ‘qualifying’ biomarkers in I-SPY2). We then further filtered this list to include only biomarkers that were detectable in organoid cultures, based on our assessment of the HUB database, and excluded proprietary biomarkers (e.g., BluePrint/MammaPrint (27,28)), resulting in a list of 10 out of the 23 published qualifying gene expression markers that could be assessed in organoids **(Figure 2C)**.

**Figure 2.**
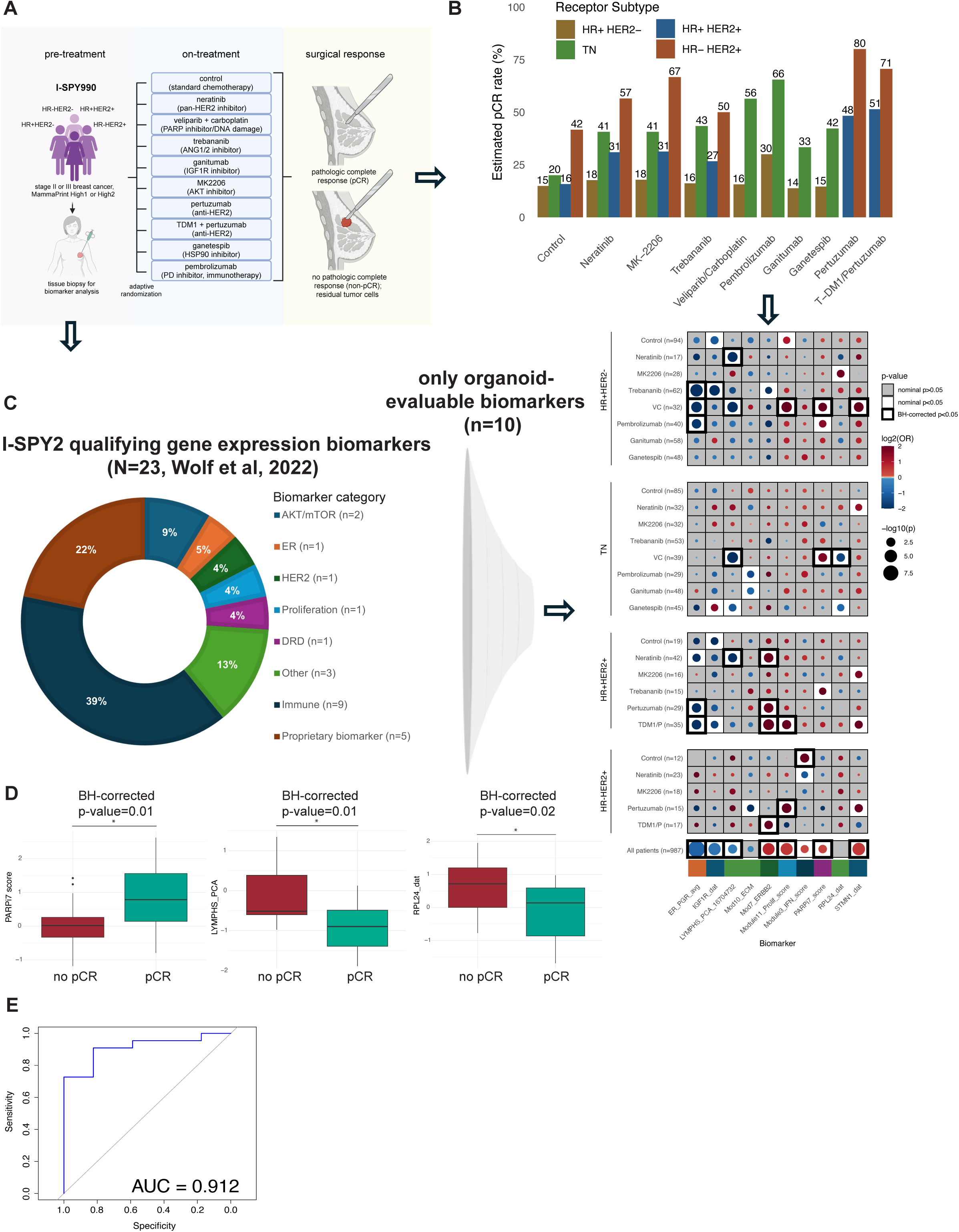
Systematic selection of I-SPY2 gene expression-based biomarkers to build a predictive model of treatment response applicable to patient-derived organoids. A. Schematic of the I-SPY2 clinical trial connecting biomarker discovery to surgical outcomes (e.g., pathologic Complete Response (pCR), which is the absence of any remaining detectable tumor after pre-operative therapy as determined by pathologic assessment of resected tissue at the time of breast surgery). B. Bar graph showing predicted pathologic complete response (pCR) rates to each of the first ten treatment arms of the ISPY2 clinical trial in each breast cancer receptor subtype. Numbers shown are the % pCR. Adapted from Wolf et al., 2022. TN = triple negative. C. Twenty-three qualifying biomarkers from I-SPY2 (summarized by biomarker category in the left panel) were filtered to include only biomarkers evaluable in organoids, based on observed expression levels in the HUB dataset and exclusion of proprietary biomarkers (n=10/23). Results of pCR association analysis in the I-SPY2 clinical trial are shown for each organoid-evaluable biomarker in each treatment arm by receptor subtype, based on gene expression and clinical data from the trial (Wolf et al., 2022) (right panel). Each row is a treatment arm. Dot color (red/blue) and intensity reflect the direction and strength of association with pCR, while dot size corresponds to the significance of the association. White background boxes indicate a nominal p-value < 0.05, and black border indicates False Discovery Rate (FDR) p-value <0.05 (Benjamini-Hochberg correction, BH). DRD = DNA repair deficiency. ER = estrogen receptor. D. Distribution of the three biomarkers that were significantly differentially expressed in triple-negative tumors in cases with or without subsequent pathologic complete response (pCR) to neoadjuvant treatment with veliparib-carboplatin (VC); from left to right: PARPi7 score, LYMPHS_PCA score, and RPL24. BH = Benjamini-Hochberg. E. Receiver operating characteristic (ROC) curve showing the performance of the final multivariable logistic regression model for predicting pathologic complete response (pCR) to veliparib-carboplatin in TNBC patients. The area under the curve (AUC) quantifies the overall discriminative ability of the model, with values closer to 1.0 indicating better performance. The model incorporating both PARPi7 score and Lymphs_PCA score demonstrates an area under the curve (AUC) of 0.912, suggesting good predictive performance for pCR in TNBC patients.

For each receptor subtype and treatment combination in ISPY2, a few of these organoid-evaluable biomarkers were significantly associated with pCR (response) or non-pCR (resistance). Our goal was to propose a robust predictive model that uses multiple organoid-detectable biomarkers to predict the probability of response to a specific treatment within a receptor subtype. We tested multivariable logistic regression models based on expression levels of the 10 selected organoid-evaluable qualifying biomarkers in each of the treatment-receptor subtype combinations for the first 10 arms of the I-SPY2 trial. After refining the models, three regression models remained that had more than one statistically significant predictor of response to a treatment within a receptor subtype: PARPi7_score and LYMPHS_PCA_16704732 were statistically significant predictors of response to veliparib-carboplatin (VC) in TNBC patients, IGF1R and LYMPHS_PCA_16704732 were statistically significant predictors of response to ganitumab in HR+HER2-patients, and Mod7_ERBB2, Module11_Prolif_score, and ER_PGR_avg were significant in a model predicting response to TDM1/P in HR+HER2+ patients.

Here, we focused on VC in TNBC patients, an investigational combination which includes the poly (ADP-ribose) polymerase (PARP) -1 and -2 inhibitor veliparib with carboplatin, a platinum chemotherapy that acts as an alkylating agent, causing cross-linking between and within DNA strands (29). Both drugs have single-agent anti-tumor activity in patients with BRCA mutation-associated breast cancer and triple-negative breast cancer, and it has been hypothesized that they could synergize when given together (30–33), though a more recent report (34) that evaluated the contributions of each of these drugs to improved long-term outcomes in TNBC demonstrated that carboplatin alone and not veliparib contributed to improved outcomes, highlighting the importance of studying resistance to platinum chemotherapies.

Three biomarkers (PARPi7, LYMPHS_PCA, and RPL24) within the set of 10 organoid-evaluable biomarkers were individually significantly associated with pCR to VC after Benjamini-Hochberg (BH) correction (**Figure 2D).** The PARPi7 gene set consists mostly of DNA repair-related genes and was positively associated with treatment response(35). RPL24 is a single-gene biomarker for the RPL24 gene that encodes the large ribosomal subunit protein eL24(36). RPL24 expression is commonly elevated in breast cancer, and depletion or structural alteration of RPL24 has been shown to significantly impair human breast cancer cell viability (37,38). LYMPHS_PCA is a gene set also mostly representing ribosomal proteins and cytoplasmic translation (39), and both were negatively associated with treatment response (5). As described above, in the multivariate logistic regression model, only PARPi7 and LYMPHS_PCA remained statistically significant. The area under the curve for the ROC curve of the model using PARPi7 and LYMPHS_PCA to predict the probability of pCR to veliparib-carboplatin in triple-negative patients was 0.912 (**Figure 2E**). We chose this model to test whether it can predict treatment response to veliparib-platinum chemotherapy (VP) in triple-negative organoid cultures based on organoid gene expression.

### TNBC Organoids (TORG) recapitulate subtypes of basal breast cancers and express key breast cancer gene signatures

To further evaluate TNBC subtypes and test the feasibility of predicting treatment response in organoids and modeling drug sensitivity and resistance states in organoids, we established a second, larger TNBC breast tumor organoid biobank (termed TNBC TORG) to expand the number of TNBCs in our dataset **(Figure 3A)**. Following previously published organoid culturing methods (40), this TNBC TORG biobank consists of 18 organoid cultures derived from primary triple-negative breast tumors and was further enriched for inflammatory breast cancer (IBC; n=8) and metaplastic cancers (n=2), aggressive forms of breast cancer that are associated with treatment resistance (**Table 2)**. Most cancers were grade 3 (78%) and clinically resistant to prior neoadjuvant chemotherapy (78%). Ki67 expression was evident in all cases that underwent additional immunohistochemical staining (n=10) with a range of expression levels, consistent with clinical variation in high-grade features. We observed various organoid morphologies across the biobank, including botryoid, solid, cystic, and metaplastic/spindle cell phenotypes (**Figure 3B**).

**Figure 3.**
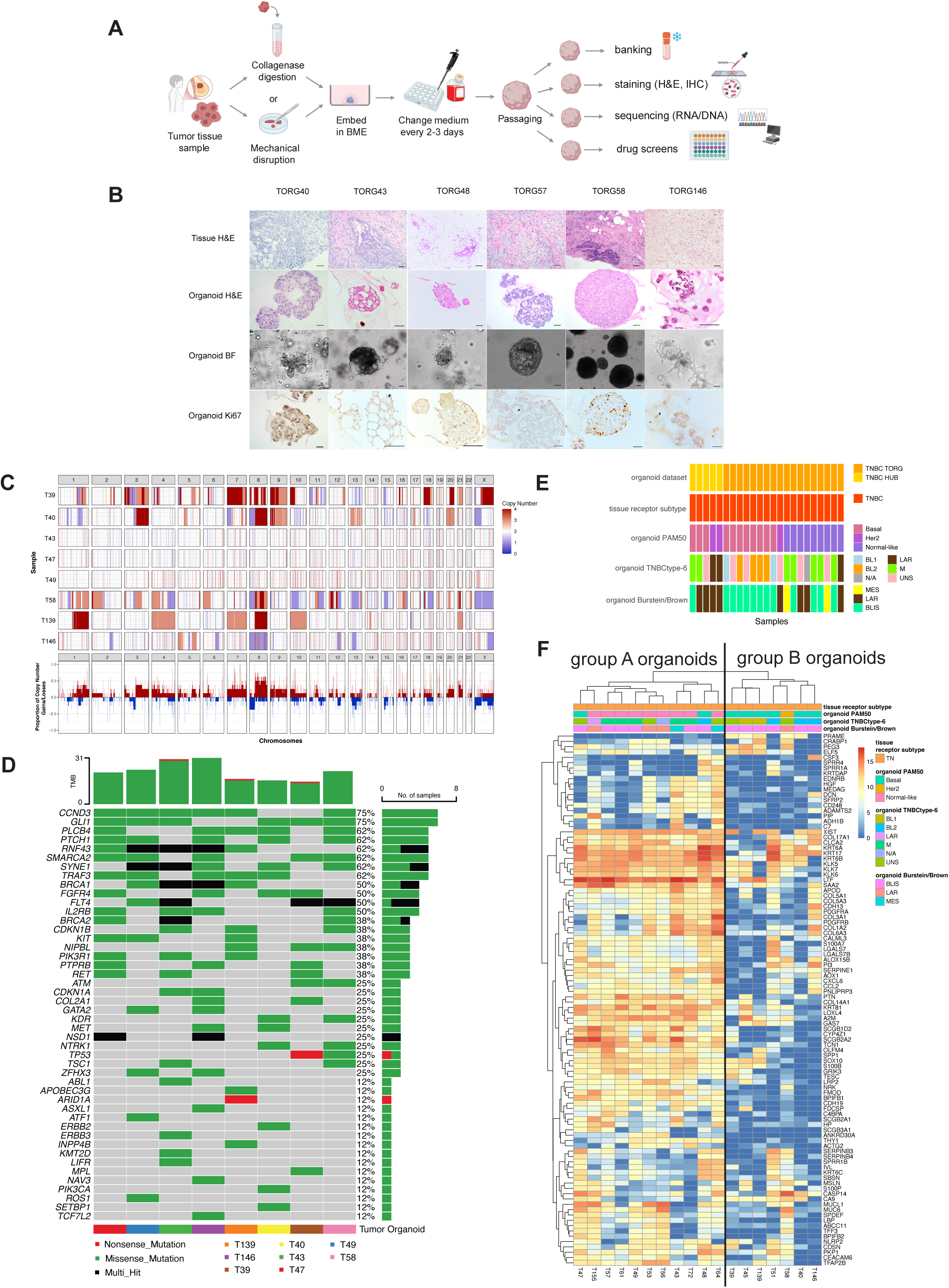
Generation and characterization of a new TNBC organoid bank comprising multiple TNBC subtypes. A. Tissue processing and organoid culturing pipeline for establishing organoids from surgical specimens. BME = basement membrane extract. H&E = hematoxylin & eosin. IHC = immunohistochemistry. B. Representative organoid cultures show diverse morphologies and phenotypes in culture, including botryoid, solid, cystic, and metaplastic/spindle cell phenotypes. For each case, histology of the tumor tissue and embedded derived organoids (stained by hematoxylin and eosin (H&E)), brightfield (BF) image of the organoids in culture, and results of immunohistochemical staining for Ki67 are shown. Scale bar = 100 μm. C. Genomic copy number variation (CNV) profiles were assessed for the indicated breast cancer organoids (n=8) using whole-genome sequencing. Each row represents an individual organoid culture, and each column represents a genomic region ordered by chromosomal position. Gains (amplifications) and losses (deletions) are indicated by red and blue colors, respectively. Quantification of recurrent gains and losses across the dataset is shown in the last row. D. Summary plot showing single-nucleotide variants (SNVs) and insertions/deletions in a representative subset of the tumor organoids (n=8), revealing the presence of mutations in breast cancer-associated genes across the biobank. E. TNBC TORG biobank reflects diverse breast cancer subtypes, identified by both general and TNBC-specific classifications. Subtyping of HUB TNBC and TNBC TORG organoids based on gene expression is shown for PAM50, as well as TNBC-specific subtypes TNBCtype-6 and Burstein/Brown subtypes, compared to the receptor subtype of the original tumor specimen. TNBC = triple negative breast cancer. BL1/BL2 = basal-like 1/2, BLIS = basal-like immunosuppressed, LAR = luminal androgen receptor, M/MES = mesenchymal, UNS = a subtype could not be selected by the algorithm, N/A = not available (sample did not pass the estrogen receptor (ER) expression filter). F. Unsupervised clustering of TNBC TORG organoids based on the top 100 most differentially expressed genes across the dataset identifies two subclusters, A and B. Annotations for each of the indicated subtype classification schemas are shown. TN = triple negative. BL1/BL2 = basal-like 1/2, BLIS = basal-like immunosuppressed, LAR = luminal androgen receptor, M/MES = mesenchymal, UNS = a subtype could not be selected by the algorithm, N/A = not available (sample did not pass the estrogen receptor (ER) expression filter).

**Table 2.** Summary of clinical and organoid culture characteristics of TNBC TORG included in the gene expression analysis (n=18).

|  |  |  | Triple-Negative (n=18) |
| --- | --- | --- | --- |
| Clinical characteristics | Grade | 1 | 1 (5.5%) |
|  |  | 2 | 3 (16.5%) |
|  |  | 3 | 14 (78%) |
|  | Tumor histological type | Ductal carcinoma | 16 (89%) |
|  |  | Lobular carcinoma | 0 |
|  |  | Other | 2 (11%) |
|  | Inflammatory breast cancer | Yes | 8 (44%) |
|  |  | No | 10 (56%) |
|  | Neoadjuvant therapy | None | 4 (22%) |
|  |  | Chemotherapy | 14 (78%) |
| Organoid characteristics | Medium type | Type 1 | 13 (72%) |
|  |  | Type 2 | 5 (28%) |
|  | Organoid morphology | Solid | 13 (72%) |
|  |  | Mixed | 5 (28%) |

CNV analysis revealed that most TNBC TORGs (5/8 tested cases; 62.5%) were heavily enriched for copy number variations (**Figure 3C**), including those generally associated with breast cancers, including amplification of chromosomes 1q and 8q (41). Within TNBCs, some subtypes are associated with high genomic instability, whereas others have a relatively stable genome, which is also reflected in our findings (42,43). Analysis of single-nucleotide variants (SNVs) and insertions/deletions in this subset of 8 tumor organoids representing organoids in both subgroups A and B and most TNBC subtypes confirmed the presence of mutations in breast cancer-associated genes across the biobank (**Figure 3D)**.

TNBC TORG gene expression data were normalized and integrated with triple-negative TCGA breast cancer tissues in the same way as the HUB biobank. We confirmed that these TNBC TORG similarly showed variable expression levels of the 50 Hallmark and 50 I-SPY2 compendium biomarkers. HER2 biomarker scores were highly variable, which suggests that some of these TNBC organoids might express HER2 even if the original cases did not meet the clinical criteria for HER2-positivity. This is interesting from a treatment perspective, as it is now known that HER2-low cancers are a targetable subset of TNBCs (44,45).

We used this expanded TNBC TORG dataset with the TNBC HUB organoids to further assess TNBC subtyping schemas in organoid cultures (**Figure 3E**). The most frequent TNBC-type-6 classifications were basal-like subtypes BL1 and BL2 (26% combined) and mesenchymal (M; 30%) (46). No immunomodulatory (IM) and mesenchymal stem-like (MSL) samples were identified in our biobank, which further supports the observation that led to the refined TNBCtype-4, which is that the IM and MSL TNBC subtypes represent tumors with substantial infiltrating lymphocytes and tumor-associated stromal cells, respectively, that are not preserved in organoids (46). Using Burstein/Brown subtypes (47), basal-like immunosuppressed (BLIS) was the most frequent subtype among the TNBC organoids (57%), with no basal-like immune-activated (BLIA) cases, again as expected for organoids.

Unsupervised clustering based on the top 100 most differentially expressed genes divided the TNBC TORG organoids into two subclusters (A, B) (**Figure 3F**). Correlating the clusters with the previous subtyping results, the organoids in group B generally had higher basal-like scores, histology associated with higher grade, a markedly faster growth rate in culture, and more extensive chromosomal alterations compared to organoids in group A. Group A organoids generally did not have as many copy number alterations, but did exhibit mutations in breast cancer-associated genes, including *PTEN*, *TP53*, and *PALB2*, and expressed Hallmark gene signatures, including higher Hedgehog signaling and estrogen/androgen response signatures compared with organoids in group B (BH-corrected p-value <0.05).

### Validation of a clinical predictive model for treatment response using TNBC organoids in vitro

Using the gene expression biomarker data from our second, larger TNBC TORG biobank (n=18), we tested the here developed model for predicting organoid treatment response to veliparib-platinum chemotherapy (VP). We ranked the eighteen organoids from the highest predicted sensitivity score to the lowest predicted sensitivity score (**Figure 4A**), with the HUB TNBC organoids, the TNBC TCGA tissues, and the I-SPY990 TNBC tissues as a reference. TNBC TORGs, TNBC I-SPY tissues, and TNBC TCGA tissues spanned the full range from predicted high VP sensitivity to high VP resistance, whereas the smaller set of TNBC HUB organoids showed a trend toward lower predicted sensitivity to VP.

**Figure 4.**
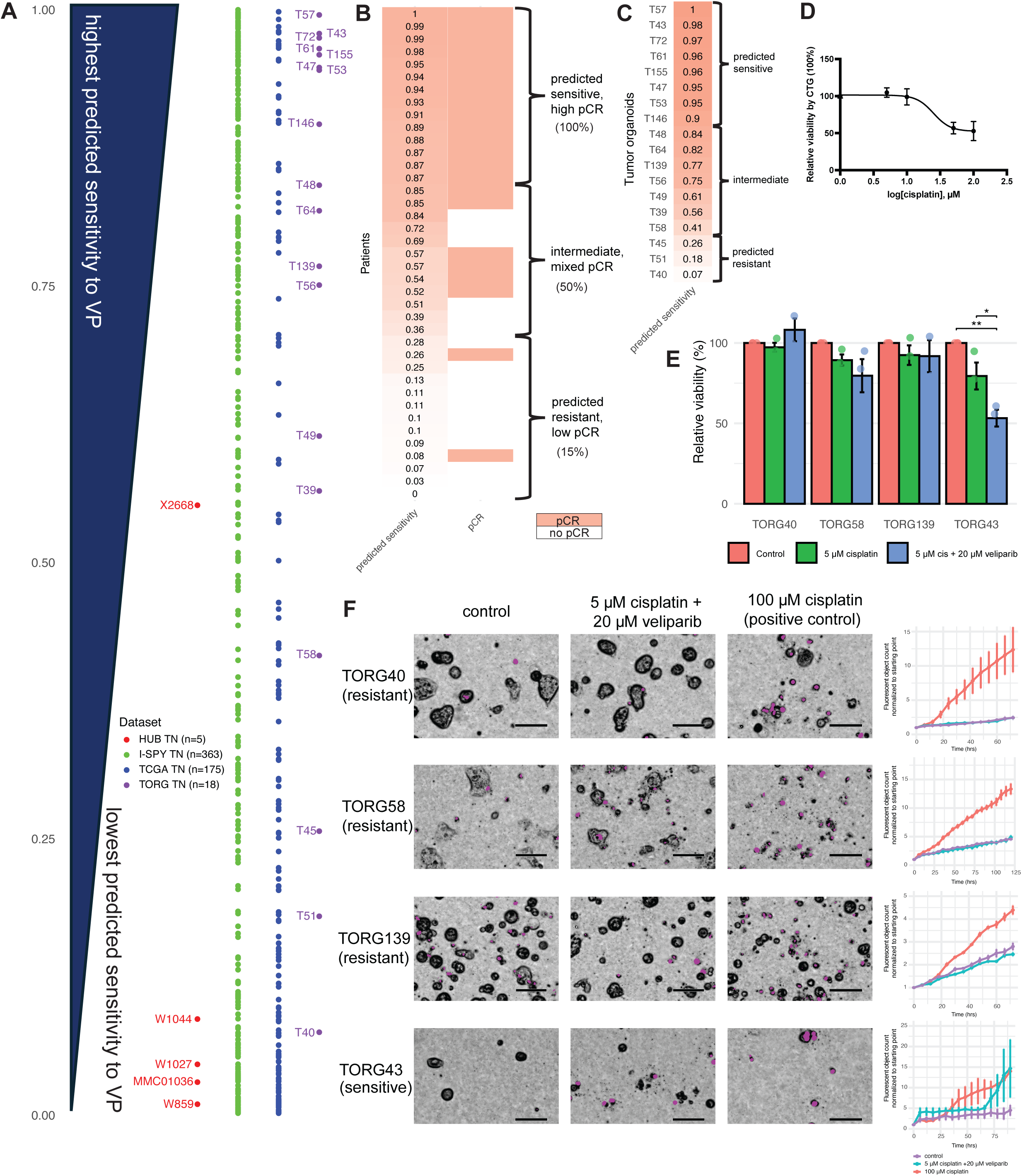
In vitro evaluation of the predictive model of treatment response. A. Ranking of organoids from the highest predicted sensitivity score to the lowest predicted sensitivity score using the predictive model of treatment response to veliparib-platinum chemotherapy (VP) in TNBC, as compared to triple-negative I-SPY2 and triple-negative TCGA tumors, shows that TNBC TORG span the full range of predicted sensitivity to VP. TN = triple negative. TCGA = The Cancer Genome Atlas. B. Ranking of triple-negative I-SPY2 patients treated with VP by predicted sensitivity scores to VP using the predictive model of treatment response to veliparib-platinum chemotherapy (VP) in TNBC, and division in tertiles distinguish a sensitive (n=14/39; 36%), intermediate (n=12/39; 31%), and resistant group (n=13/39; 33%), as confirmed by pathologic complete response (pCR) outcomes (pCR rates 100%, 50%, and 15%, respectively). C. Ranking of predicted sensitivity scores to VP in TNBC TORG using the predictive model of treatment response to veliparib-platinum chemotherapy (VP) in TNBC, also divides the organoids into a predicted sensitive (n=8/18; 44%), predicted intermediate (n=7/18; 39%), and predicted resistant group (n=3/18; 17%) based on the threshold values identified in the I-SPY2 VP-TNBC dataset. D. In vitro dose-response curve in TORG40 demonstrates resistance to cisplatin. Relative cell viability (%) was determined at T=72 hours using a luminescence-based cell viability assay (Cell-Titer-Glo). Data are expressed as the mean ± standard error of the mean (SEM). E. In vitro evaluation of the predicted response to veliparib-platinum chemotherapy (VP) by the mathematical model, in a predicted resistant (TORG40), two predicted intermediate (TORG58 and TORG139), and one predicted sensitive organoid (TORG43), using a luminescence-based cell viability assay (Cell-Titer-Glo), shows results concordant with response predictions. Bars show mean relative cell viability (%) compared to untreated control, with error bars showing the standard error of the mean (SEM) for three independent experiments. Differences between treatment conditions were tested using the ANOVA test for each organoid culture, with a post-hoc Tukey test. **= p-value<0.01, *= p-value<0.05. cis = cisplatin. F. Live-cell imaging results corresponding with the experiments in Figure 5E illustrate the lack of drug response (lack of cell death) after 72-120 hours of drug treatment with 5 μM cisplatin + 20 μM veliparib in TORG40 (predicted resistant), TORG58 (predicted intermediate), and TORG139 (predicted intermediate), while treatment with 5 μM cisplatin + 20 μM veliparib induces significant cell death in TORG43 (predicted sensitive), representative of three independent experiments. Purple color indicates fluorescence from a cell-death dye (CellTox), as captured by the Incucyte, with quantification in the right panels. 100 μM cisplatin was included as a positive control. Data are expressed as the mean ± standard error of the mean (SEM). Statistical testing (Kruskal-Wallis test) was not significant. Scale bar = 100 μm.

We evaluated the calculated sensitivity scores for I-SPY patients using our predictive model and correlated them with pCR (**Figure 4B**). We identified one tertile of patients who had high predicted sensitivity scores (>0.85) and all achieved a pCR (n=14/14 pCR, 100%), one tertile of patients with low predicted sensitivity scores (<0.3) and who mostly did not achieve a pCR (n=2/13 pCR, 15%), and an intermediate subgroup with predicted sensitivity scores between 0.3-0.85 who had mixed pCR outcomes (n=6/12 pCR, 50%). Similarly, translating these thresholds to the TORGs set, we identified a predicted sensitive group (n=8), an intermediate group (n=7), and a predicted resistant group (n=3) (**Figure 4C**).

To evaluate the response prediction, we tested four cultures in total: 1 predicted sensitive culture (TORG43) and 1 predicted resistant culture (TORG40), and two intermediate cultures (TORG58, TORG139). Based on a dose-response curve of cisplatin in TORG40, the most predicted resistant organoid, we selected 5 μM as the fixed low-dose cisplatin concentration for any combination treatment (**Figure 4D**). After 72 hours of drug treatment for fast-growing organoids (TORG40, TORG139), and 96-120 hours for moderate to slow-growing organoids (TORG43, TORG58), with cisplatin monotherapy (5 μM), or cisplatin combination therapy with veliparib in two doses (20 μM veliparib and 10 μM veliparib, respectively), cell viability was assessed using Cell-Titer-Glo and live-cell imaging (**Figure 4E-F**). Predicted-resistant/intermediate organoids TORG40, TORG58, and TORG139 were highly resistant to cisplatin monotherapy and VP combination therapy, while predicted-sensitive organoid TORG43 was sensitive to the high dose veliparib in combination with cisplatin (5 μM cisplatin + 20 μM veliparib). Altogether, response predictions by the biomarker-based model in vitro were successfully validated in the selected organoids.

### Drug screen in TNBC organoids identifies strategies to overcome resistance to cisplatin

High-throughput screening with a fixed-dose backbone therapy (“anchor” drugs) is useful for identifying combination therapies(48). With the aim of finding alternative treatment strategies to overcome resistance to platinum chemotherapy, we selected TORG40, a robust culture with the highest predicted and subsequently validated resistance to VP, and performed a high-throughput drug screen of 386 small-molecule inhibitors at 10 doses in the presence and absence of a fixed concentration of cisplatin to find synergistic or additive small molecule inhibitors with cisplatin (median Z’ factor of the screen was 0.788; interquartile range 0.246-0.837(49)). Cisplatin and carboplatin have highly similar mechanisms of action, with highly correlated sensitivity/resistance profiles in vitro (50). Clinically, cisplatin and carboplatin are both used to treat breast cancer and have similar efficacy, while carboplatin reportedly has less toxicity than cisplatin (51). For in vitro drug screens, we chose cisplatin, given its greater chemical stability and higher potency than carboplatin (51).

To identify the top hits in our drug screen, we first ranked the 386 drugs by largest difference in area under the curve (AUC) of the dose-response curve of the drug without cisplatin compared with the drug with cisplatin (**Figure 5A**). We found that for 20/386 (5.2%) of tested compounds, the AUC with cisplatin decreased by 40% or more compared with the monotherapy, indicating an increased efficacy with cisplatin. 76/386 (19.7%) had a decrease in AUC between 20-40%, and the remaining 290/386 (75.1%) had less than 10% to no change in AUC, or an increase in AUC with cisplatin, indicating no important improvement of efficacy with the treatment combination compared with the monotherapy. The difference in AUCs between the condition without cisplatin and with cisplatin was used to rank the compounds and select the top 5% of compounds in the screen with the largest reduction in AUC with the addition of 5 μM cisplatin (n=19). 17/19 (89%) compounds were also among the top 20% of compounds with the smallest IC50 value when given in combination with cisplatin. From these, we selected 8 compounds with various mechanisms of action (HDAC inhibitors (epigenetics), Bcl-2 inhibitors (apoptosis), an inhibitor of DNA synthesis (DNA damage repair), an IAP inhibitor (apoptosis), and an aurora kinase inhibitor (cell cycle regulation) with a maximum of 3 compounds per drug target (individual dose-response curves shown for each drug, with and without cisplatin, in the original drug screen in **Figure 5B**). We validated these compounds with and without the addition of 5 μM cisplatin in the same organoid culture (TORG40) in the Incucyte live-cell analysis system using a secondary, CellTox™ Green Cytotoxicity Assay, which is a fluorescence and imaging-based assay. We confirmed an additive effect of cisplatin with all tested compounds at a minimum of one of the concentration levels tested, as quantified by an increase in total normalized fluorescent object count in the combination treatment with cisplatin versus the monotherapy, along with matching morphological changes by Brightfield imaging (**Figure 5C**).

**Figure 5.**
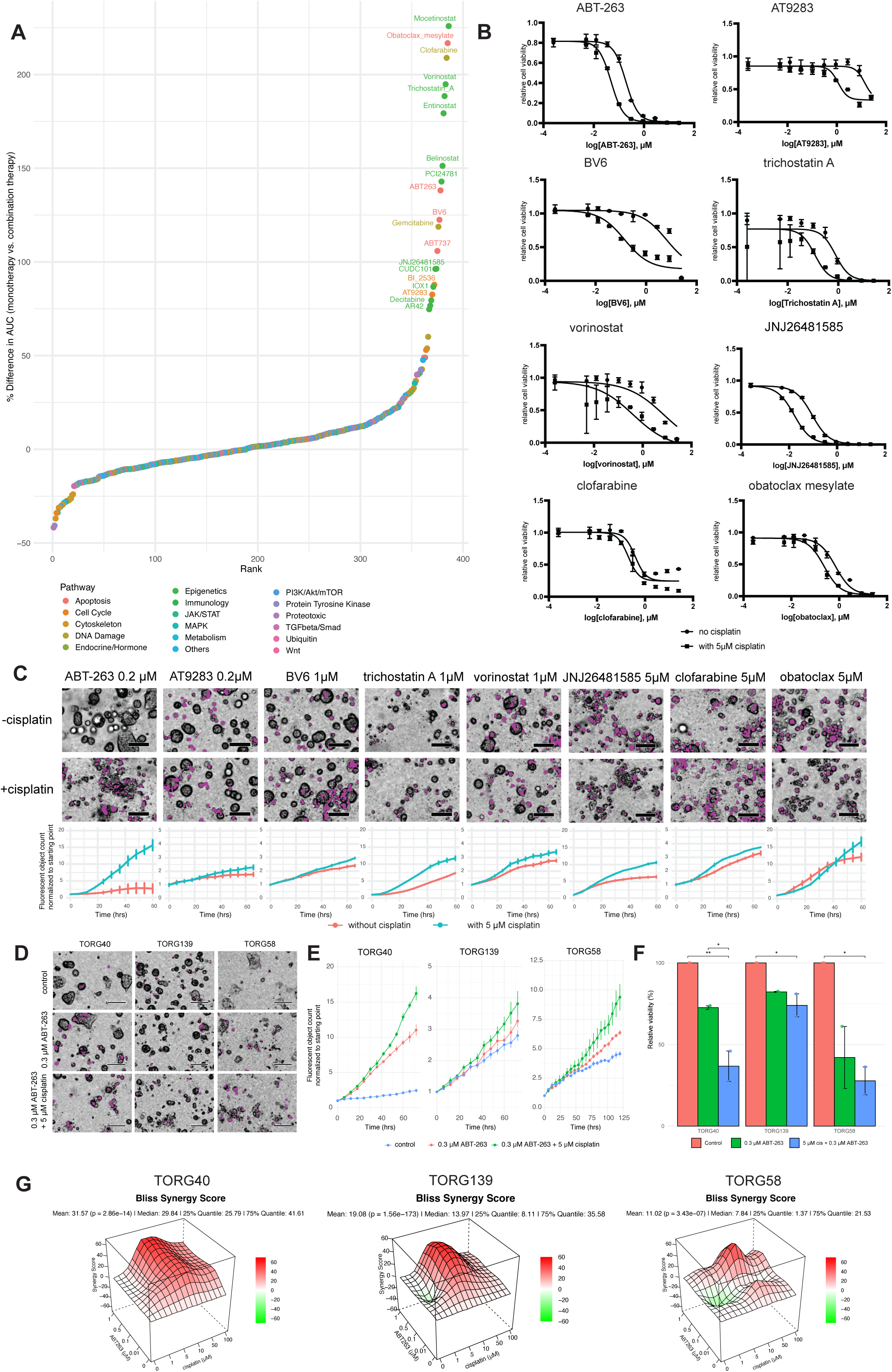
A high-throughput drug screen in a tumor organoid resistant to VP reveals new treatment strategies to overcome resistance to platinum chemotherapy. A. Ranking of 386 drugs in high-throughput drug screen by the percent difference in area under the curve (AUC) of the dose-response curve of the drug compound without cisplatin (monotherapy) compared with the AUC of the drug with cisplatin (combination therapy). Each dot represents one drug in the drug screen, sorted by increasing rank number, with the highest rank = the highest difference. The top 5% of compounds with the largest reduction in AUC with cisplatin are labeled with the drug name. The colors of the dots correspond with the mechanism of action of the drug. B. Average drug dose-response curves for a selection of the top combination therapies in the 386-compound drug screen (squares indicating the average dose-response curve for the drug + 5 μM cisplatin, and circles indicating the average for the drug as a monotherapy) in TORG40 determined at T=72 hours using a luminescence-based cell viability assay (Cell-Titer-Glo). Data are expressed as the mean ± standard error of the mean (SEM). C. Validation of the top combination treatments from the drug screen in TORG40 using a secondary, imaging-based assay in TORG40 captured in the Incucyte. Purple color indicates fluorescence from a cell-death dye (CellTox). Graphs show total fluorescent object count at the final timepoint (T=60-66 hours) normalized to the starting point (T=0), with the blue lines indicating the average normalized fluorescent object count for the drug + 5 μM cisplatin, and the pink lines indicating the average for the drug as a monotherapy. Data are expressed as the mean ± standard error of the mean (SEM). Scale bar = 100 μm. Statistical testing (Wilcoxon test) was not significant. D. Validation of the cisplatin + Bcl-2 inhibitor treatment (ABT-263) condition in two other in vitro resistant cultures to veliparib-platinum chemotherapy (VP) by brightfield and fluorescent imaging show superior drug response with 5 μM cisplatin + 0.3 μM ABT-263 (bottom) compared with 0.3 μM ABT-263 alone (middle) after 72 hours of drug treatment in TORG40 and TORG139, and 120 hours in TORG58. Purple color indicates fluorescence from a cell-death dye (CellTox), as captured by the Incucyte. Scale bar = 100 μm. E. Cell death curves corresponding with images in Figure 5D, showing the effects of Bcl-2 inhibitor treatment (ABT-263) with and without cisplatin as evaluated at 72 hours for TORG40 and TORG139, and 120 hours for TORG58 by live-cell imaging and a fluorescence assay (CellTox; total fluorescent object count at endpoint (72 or 120 hours) normalized to the starting point (T=0)). Fluorescence data are expressed as the mean ± standard error of the mean (SEM). Statistical testing (Kruskal-Wallis test) was not significant. F. Validation of the cisplatin + Bcl-2 inhibitor treatment combination in two other in vitro resistant cultures to veliparib-platinum chemotherapy (VP) using a cell viability assay (Cell-Titer-Glo) shows an additive effect of cisplatin to ABT-263 measured at 72 hours for TORG40 and TORG139, and 120 hours for TORG58. Bars show mean relative cell viability (%), compared to the untreated control ± standard error of the mean (SEM). Differences between treatment conditions were tested using the ANOVA test for each organoid culture, with a post-hoc Tukey test. **= p-value<0.01, *= p-value<0.05. cis = cisplatin. G. 3D plots showing the Bliss synergy analysis results for ABT-263 with cisplatin for TORG40, TORG139, and TORG58. The organoids have mean Bliss synergy scores >10 (p-value <0.05), suggesting probable synergy between ABT-263 and cisplatin, particularly at 5-10 μM cisplatin doses, representative of two independent experiments.

Lastly, we sought to validate the clinically most promising drug combination of cisplatin + a Bcl-2 inhibitor (ABT-263) in other resistant organoid cultures. In TORG58 and TORG139, two other in vitro validated VP-resistant organoids, a consistently increased efficacy of ABT-263 + cisplatin over ABT-263 monotherapy was observed, measured as a decrease in viable organoid morphology by brightfield (**Figure 5D)**, increased cell death by CellTox (**Figure 5E)**, and decreased cell viability by Cell-Titer-Glo (**Figure 5F)**. This was consistent with the original screen in TORG40. We performed formal synergy experiments in all three organoids by testing increasing doses of cisplatin in combination with increasing doses of ABT-263 to identify dose combinations where a synergistic effect could be observed. Bliss synergy analysis results revealed mean Bliss synergy scores between 11.02 and 31.57 (p<0.05), with the strongest synergy observed at 5-10 μM cisplatin doses, suggesting probable synergy between ABT-263 and cisplatin(52). These results highlight Bcl-2 as a promising drug target across multiple models of resistance to VP.

## Discussion

Here, we describe the first application of a predictive model obtained from clinical patient biomarker data to predict treatment response or resistance in organoids. We used this model to select biomarker-matched treatment-resistant organoid cultures from a biobank and tested alternative treatment strategies. This proof-of-principle study demonstrates the ability of breast cancer organoids to bridge the gap between correlative clinical science and experimental biology. While studies have shown concordance in expression of focused sets of biomarkers between patient-derived organoids (PDOs) and patient-derived xenografts (PDXs) (6,7,53), experimental validations of patient tissue-derived biomarkers in preclinical models are still scarce (54). With the immense rise of genomics studies and increasing numbers of predictive biomarkers, there is a need to translate in silico transcriptomic findings to functional in vitro tools that can then be used to test or stratify large numbers of investigational therapies.

To this end, we utilized gene expression from an existing organoid biobank and developed a new set of TNBC organoids, derived from both pre-treatment and post-treatment tissues, covering a wide range of histological subtypes and including rare subtypes that are associated with treatment resistance that are often overlooked, such as metaplastic and inflammatory breast cancers. This also includes a set of organoids that classify as “normal-type” by PAM50. We hypothesize that the “normal-type” classification could be explained by either LAR tumors or by heterogeneity of the original tumor tissue, where admixed populations of cells may not be represented well by a bulk analysis technique and a categorical classification. Interestingly, another subset of TNBC organoids was particularly aggressive in culture, as characterized by fast growth in vitro, high expression of MYC and proliferation signatures, and more extensive genomic copy number aberrations. They represent a subgroup of breast cancers in exceptionally high need of adequate treatment.

We modeled organoid resistance states using qualifying gene expression-based biomarkers and clinical data from the I-SPY2 trial. These qualifying biomarkers were pre-specified for prospective testing in I-SPY2 for their specific response-predictive value based on existing evidence suggesting a role in treatment response prediction (5), making them especially interesting and meaningful to evaluate in organoid models. The most significant model using multiple biomarkers that are expressed in our organoid models and correlating with clinical outcomes in TNBC patients in the I-SPY2 trial was a model that predicted response to veliparib-platinum chemotherapy. This is a particularly clinically relevant model since platinum chemotherapies are often prescribed to patients with triple-negative breast cancer (55–58), but non-response is frequent, and new approaches to overcome platinum-chemotherapy resistance are needed. Moreover, this model exemplifies that DNA damage genes (in the PARPi7 signature) can predict response to chemotherapy and/or PARP inhibition at the gene expression level, not only in patient tissues (5), but also in organoids. The organoids with the highest predicted and validated resistance to platinum chemotherapy plus veliparib were mostly basal-like organoids (basal by PAM50 subtype, BL1 or BL2 by TNBC-type and BLIS by Burstein/Brown subtypes), suggesting that findings from our drug screen may be most applicable to patients with basal-like tumors.

Multiple clinical studies, including I-SPY, have established platinum therapy as part of the standard of care for the treatment of TNBC. A phase II study randomizing patients with early-stage TNBC to preoperative cisplatin versus paclitaxel found that the Residual Cancer Burden (RCB)-0/1 rate – indicating pathologic complete response or minimal residual cancer burden – was 26.4% with cisplatin and 22.3% with paclitaxel (59). Other phase II studies exclusively for patients with *BRCA1* mutation demonstrated markedly higher pCR rates from 61% to 90% after neoadjuvant cisplatin monotherapy (60). In I-SPY2, the response to veliparib-carboplatin followed by doxorubicin/cyclophosphamide chemotherapy in triple-negative patients was 56% (5). The phase III BrighTNess trial showed that adding carboplatin to standard neoadjuvant chemotherapy for stage II-III TNBC improved event-free survival, whereas adding veliparib did not impact long-term outcomes after 4.5 years of follow-up (34). Thus, we chose to focus our drug screen on overcoming resistance to platinum chemotherapy. Using our predictive model, our TNBC TORGs displayed a wide range of predicted sensitivity and resistance to VC, covering the full range of sensitivity scores in both I-SPY and TCGA samples. Our subsequent drug screen identified HDAC inhibitors, Bcl-2 inhibitors, and IAP inhibitors to be most synergistic with cisplatin, and the Bcl-2 inhibitor ABT-263 was validated in two other organoid models with demonstrated resistance to VC.

Interestingly, this drug had previously displayed synergy with oxaliplatin in a colon cancer organoid drug screen (9). Other studies found that Bcl-2 inhibitors can also synergize with PARP inhibitors in ovarian cancer, and that high levels of Bcl-2 mediate cisplatin resistance in bladder cancers (61,62). In breast cancer, the efficacy of Bcl-2 inhibitors has been tested with endocrine therapies with mixed results (63,64), though they have been proven to be effective and approved by the FDA for the treatment of certain hematological cancers, including chronic lymphocytic leukemia (CLL) and acute myeloid leukemia (AML) (65,66). Ongoing clinical trials in breast cancer are assessing combination therapies of Bcl-2 inhibitors with PARP inhibitors and other novel agents, but not yet with platinum chemotherapies (67,68). Our study suggests this strategy should be tested further to overcome platinum resistance in aggressive triple-negative breast cancer subtypes.

A general limitation of our study is that current culture methods applied in this research to create and maintain tumor organoid cultures were not yet suited to preserve the immune environment, limiting us to a subset of clinical biomarkers that are evaluable in organoid cultures and, therefore, limiting the treatment mechanisms/arms we can model. Given the success that immune checkpoint inhibition has shown for certain patients with TNBC and the expected role of immunotherapies in the neoadjuvant treatment for TNBC (69), it would be valuable to test our approach in other organoid models incorporating immune cells in culture, such as early-stage organoids and air-liquid interface models (70–73), and organoids with other culture media that are optimized to support various components of the tumor microenvironment in culture, to better model the interactions between the tumor and its immune environment.

Secondly, our study utilized bulk gene expression and thus did not evaluate the heterogeneity inherent within a single tumor and its relationship to the complex composition of organoid culture, which we observed in our dataset. Contributions from different subclones or subpopulations of tumor cells could not be delineated, and signals from tumor cells might be diluted by the presence of normal or stromal cells present in the culture, particularly during early passages. With the I-SPY2 trial continuously enrolling new patients, the clinical and biomarker data resource will continue to grow, enabling the continuous improvement of predictive models. Moreover, we are currently working on establishing I-SPY organoids from patients enrolled in the I-SPY clinical trial, to benchmark organoid data and characteristics against the original pre- and post-treatment tissue samples of patients.

In summary, by combining computational analyses of large molecular and clinical datasets with organoid model systems, this proof-of-principle study demonstrated the utility of matching I-SPY2 resistance biomarkers and signatures to residual disease tumor organoid cultures. This research demonstrated that tumor organoid cultures can be utilized in the laboratory to mimic drug resistance states and serve as a valuable complement to existing research models of breast cancer for preclinical drug testing informed by tumor biology. Ultimately, this approach can be used to reveal new therapeutic vulnerabilities and resistance mechanisms, enabling faster and more successful translational studies and increasing treatment options for resistant diseases.

## Acknowledgments

The authors thank Hans Clevers for 293T-Noggin and L-Wnt3a cell lines, and Calvin Kuo for 293T-RSPO1, and Joan Brugge and Laura Selfors for their helpful discussions and advice. We thank Mackenzie Boedicker, Griffin Boedicker, Alexis Cook, and Pravin Phadatare for technical assistance, the Inflammatory Breast Cancer Program at Dana-Farber Cancer Institute, and the DF/HCC Breast SPORE: Specialized Program of Research Excellence (SPORE) (NCI 1P50CA168504), the UCSF Breast Care Center surgical teams including Jasmine Wong and the Breast Care Center interns for assistance with tissue specimen. We further thank the ICCB-Longwood Screening Facility at Harvard Medical School and the Wynton HPC Co-Op cluster, which is supported by UCSF research faculty and UCSF institutional funds. Funding was from the Susan G. Komen Foundation and METAvivor (to J.M.R.), and Breast Cancer Foundation (to L.V.V. and J.M.R.). T.B.V.B. was supported by the ROSANNA Fund, Hendrik Muller Fund, Netherland-America Foundation, and the Foundation of Renswoude. J.M.R. is a Chan-Zuckerberg Biohub – San Francisco investigator. The content is solely the responsibility of the authors and does not necessarily represent the official views of the NIH/NCI.

## Author contributions

Conceptualization: JMR, TBVB, LvV, DW

Data curation: TBVB, DW, IN, MCB, JL

Data analysis: TBVB, DW

Funding acquisition: JMR

Investigation: TBVB, JMR, DW, KM, MCB, SW, AP, SC, JL

Methodology: TBVB, JMR, DW, IH

Project administration: TBVB, JMR

Software: IH, DW

Sample acquisition/Resources: JMR, DD, BO, FL, IN, DW, LE, LvV

Supervision: JMR, LvV, BMTB

Validation: TBVB, DW, MCB, SC, JL

Visualization: TBVB, DW, JL, SC

Writing – original draft: TBVB, MCB, JMR

Writing – review & editing: all authors

## Methods

### **I-** SPY2 trial biomarker resource to develop predictive models of treatment response

Matched biomarker and clinical outcomes data of patients of the first ten treatment arms of the I-SPY2 trial (n=986) were used to develop a mathematical prediction model for treatment response, based on the expression of biomarkers derived from patient tissue molecular data. We included in our analysis only those biomarkers that were detectable in organoid cultures and excluded proprietary biomarkers (e.g., BluePrint/MammaPrint (27,28)), resulting in a final list of 10 qualifying gene expression markers from Wolf et al. (2022) that could be assessed in organoids.

Using the I-SPY990 patient outcomes data (pathologic complete response; pCR) and the I-SPY990 patient tumor tissue expression level of these 10 organoid-evaluable biomarkers, we performed a multivariable logistic regression analysis across all receptor subgroup-treatment arm combinations. Receiver operating characteristic (ROC) plots were made using pROC in R. We selected the model that predicted response to VC in TNBC patients for further validation. Applying the predictive model of response to VC in TNBC organoids of the HUB and TNBC TORG biobanks, as well as patient tissues from the TCGA and I-SPY datasets, yielded a ranking of samples from most predicted sensitive to most predicted resistant.

### Breast Cancer Tissue Processing and Patient-Derived Organoid Culture

Tumor organoids were grown in three-dimensional cultures following a previously published protocol (40). In short, tissue samples were processed on the day of surgery. Upon arrival in the lab, tissues were photographed for documentation and, if big enough, divided into two or more 10-15 mm^3^ pieces. Of each tissue piece, if big enough, a small section was cut off and formalin-fixed and paraffin-embedded (FFPE) for histopathological analysis. In the case of multiple tissue pieces, one or more pieces of the fresh tissue were viably frozen in 90% Fetal Bovine Serum (FBS; Cytiva, Cat. No. SH30910.02) and 10% DMSO and banked in liquid nitrogen for the subsequent generation of organoid cultures. The remaining tissue piece was minced and either directly embedded in a 50 μl drop of basement membrane extract (BME) type 2 (Cultrex, Cat. No. 3532-005-02) in the center of a well in a pre-warmed 24-well tissue culture plate (Greiner, Cat. No. M9312 and CytoOne, Cat. No. CC7672-7524) or first digested by placing the minced tissue into a 50 ml conical tube containing 1 mg/ml collagenase (Sigma, Cat. No. C9407) and 20 ml AdDF+++ (Advanced DMEM/F12 (Thermo Fisher, Cat. No. 12634-028) containing 1× Glutamax (Thermo Fisher, Cat. No. 35050-061), 10mM HEPES (Thermo Fisher, Cat. No. 15630-080), and antibiotics: penicillin-streptomycin (Thermo Fisher, Cat. No. 15140-122) and Primocin (Invivogen, Cat. No. Ant-pm-05)). Tubes containing minced tissue and collagenase were wrapped in parafilm and placed on their side in an orbital shaker at 37 °C for 15 to 30 minutes. After digestion, 30 ml AdDF+++ and 2% FBS were added, and the organoids were pelleted. Further mechanical shearing was achieved by adding 10 ml AdDF+++ and sequentially pipetting with 10-, 5-, and 1-ml pipette tips before the final pellet of primary breast organoids was obtained and embedded in a 50 ul drop of BME. BME drops were allowed to harden for 20 minutes at 37°C, after which 500 ul of a fully defined type 1 or type 2 organoid culture medium (40) was placed over the droplet. Organoids were stably cultured in the default organoid medium (type 1, 72%), and in type 2 medium (28%) if stable growth was not achieved in type 1 (40). Plates were kept in humidified 37°C / 5% CO2 incubators at ambient O_2_, and the organoid medium was changed every 2-3 days. Organoids were passaged at different rates depending on the growth rate of the organoids, varying from once a week, to once in 4 or more weeks. Most organoids had a solid morphology and a moderate growth rate. 0.5 ml of TrypLE™ (Gibco, Cat. No. 12604013) per 2-3 wells of organoids was used to break down the organoids into smaller clumps of cells, following the addition of 10 mL AdDF+++, 0.2 ml FBS to deactivate the TrypLE and centrifugation at 300 g for 3 minutes. Organoids were re-plated in BME domes at ratios varying from 1:1 – 1:6 depending on individual organoid growth characteristics. Growth rate was categorized as “Fast” (split at least 1:2 every week), “Moderate” (split 1:2 every 1-4 weeks), or “Slow” (split 1:2 every 4 or more weeks). Established cultures with organoids were viably frozen on a regular basis and banked at different passage numbers using Recovery Cell Culture Freezing Medium (Life Technologies, Cat. No. 12648-010) for future use. Cultures were routinely tested for Mycoplasma infection.

Depending on growth rate and success, organoids were submitted for bulk RNA sequencing and, if possible, whole genome sequencing, in that order, after they had reached at least four passages.

### Organoid embedding and staining (Hematoxylin & Eosin staining, immunohistochemistry)

Organoids were embedded, sectioned, and stained as previously described in Rosenbluth et al. (74). In short, after embedding intact BME gels containing organoids in histogel and fixing them in formalin, they were paraffin-embedded, sectioned, and prepared for Hematoxylin & Eosin staining and immunohistochemistry. Immunohistochemistry was performed as published.

For H&E staining, slides were deparaffinized and rehydrated, and then stained in hematoxylin (Sigma Aldrich, Cat. No. MHS32-1L) for 3 min, followed by washing in water until it runs clear. Then destaining in 100% Ethanol (Fisher Scientific, Cat. No. C961Y21 (CS/1)) for 1 min and subsequently stained in eosin Y (Sigma Aldrich, Cat. No. SIAL-1170811000) for 10 min, followed by washing in 100% ethanol twice for 5 minutes each. After dehydration, the slides were covered with coverslips.

### Organoid RNA and DNA extraction

#### RNA Extraction

Organoids passage 4 or higher were harvested when confluent using TRIzol® reagent (ThermoFisher, Cat. No. 15596018). Cells were lysed using chloroform (SigmaAldrich, Cat. No. C2432-1L) in preparation for using the Purelink RNA Mini Kit® Protocol (ThermoFisher, cat# 12183020). Samples were eluted in 30 μL RNA-Free water through two serial elutions. Samples were stored at -80°C until submission for library preparation and sequencing.

#### DNA Extraction

Organoids passage 4 or higher were harvested when confluent by digesting them using TrypLE, followed by a wash in AdDF+++ and FBS, after which the pellet is resuspended in 500 μL FBS, pelleted again, and then snap-frozen in dry ice-cooled ethanol (Thomas, Cat. No C961Y21). Organoids were stored at -80°C until DNA was extracted using DNeasy Blood & Tissue Kit® (Qiagen, Cat. No. 69504). Samples were eluted in 30 μL RNA-free water. For lower-confluency samples where a low RNA yield was expected, the 30ul eluate was run through the spin cartridge a second time to maximize the RNA elution. Sample concentration was measured by Qubit analysis of 1 μL of sample. Samples were stored at -20°C until submission for library preparation and sequencing.

### RNA sequencing and analysis

Bulk RNA-sequencing and analysis of TNBC TORG breast cancer organoids were performed at Novogene. In short, after sample quality control, mRNA libraries were prepared using poly(A) enrichment of the mRNA. The library was checked with Qubit and real-time PCR for quantification and a bioanalyzer for size distribution detection. Illumina NovaSeq 6000 PE150 was used for high-throughput sequencing. Raw sequencing reads were filtered to remove adapter-contaminated and low-quality sequences. Clean reads were then aligned to the reference genome (Homo_sapiens_Ensemble_75). Gene expression levels were quantified using RSEM, providing transcript abundance estimates for downstream analyses(75).

To analyze breast cancer subtypes and I-SPY gene expression biomarker signatures in tumor organoid data, we log-transformed organoid bulk RNA-sequencing data and quantile-normalized them to a larger dataset of TCGA breast cancer tissues to enable analysis and interpretation of the organoid data in a larger, more robust context. Expression and clinical data from breast cancer samples in TCGA were downloaded from the PanCanAtlas Publications Supplemental Data (76). Quality controls were performed to confirm the successful merge of the data, including principal component analyses (PCA) of the tumor organoids together with all TCGA breast cancer samples based on the expression of the PAM50 genes.

The merged organoid-TCGA datasets were then subtyped by PAM50-subtype (genefu package in R) and two TNBC classification schemas (Burstein/Brown triple-negative classifications and TNBCtype-6). Hallmark gene set scores were calculated by gsva, and qualifying biomarkers as well as a set of exploratory, partly immune, biomarkers from the I-SPY2 clinical trial and other published gene signatures were scored in the merged organoid-TCGA tissue datasets (5,77–85).

### DNA sequencing and analysis

Whole-genome sequencing of TNBC TORG breast cancer organoids was performed at Novogene. In short, genomic DNA libraries were constructed and, after quality control, sequenced on the Illumina NovaSeq X Plus platform (PE150). Raw data were transformed into short-read sequences and recorded in FASTQ format. Quality control procedures filtered reads containing adapters or with low sequencing quality. High-quality reads were aligned to the reference genome to generate BAM files for subsequent analyses. Variant detection focused on germline mutations, including SNPs, InDels, and CNVs, due to the absence of matched tumor-normal samples. Germline CNVs were detected using Control-FREEC. Germline SNPs and InDels were identified using GATK HaplotypeCaller, with variant filtering performed using the GATK VariantFiltration module, followed by annotation using ANNOVAR (86–88).

### Sensitivity testing to cisplatin-veliparib

In TORG40, we performed a 5-point dose-response curve to cisplatin (SigmaAldrich, Cat. No. 232120-50MG) to identify the IC10 and IC50, to help select the dose for the platinum backbone therapy in later combination treatment testing in the organoids. Organoids were grown in 50 μL 100% BME gel domes in 24-well plates supplemented with organoid medium and grown to full confluency. One confluent well with organoids was collected and digested into small clumps using TrypLE, filtered, and resuspended in organoid medium containing 6% BME. Using a P100 multichannel pipette, 90 μL of organoid suspension was dispensed into 10-20 wells (depending on specific experiment protocols and the growth rate and confluency of the organoids) into a clear-bottom 96-well plate (Corning, Cat. No. 3610). Two days after plating, 30 μL of medium containing cisplatin in increasing doses was added to each of the wells in triplicate (final concentrations of cisplatin in the well: between 1 and 100 μM). After 72 hours, cell viability was assessed using Cell-Titer-Glo. Plates were read in a luminometer.

In vitro drug response to veliparib (SelleckChem, Cat. No. S1004) + cisplatin was tested in every organoid culture that had sufficient growth to perform the drug experiment in at least three biological replicates. After plating as described above, on day 2, the organoids were treated with either DMSO (negative control), 5 μM cisplatin in 0.9% NaCl (the IC10 concentration identified in experiments in TORG40), 5 μM cisplatin in 0.9% NaCl + 10 μM veliparib in DMSO, or 5 μM cisplatin in 0.9% NaCl + 20 μM veliparib in DMSO, all in technical triplicates. The final DMSO concentration in the wells was kept under 1%. For fast-growing organoids, drug experiments were stopped at T=72 hours, and cell viability was assessed using Cell-Titer-Glo. For slower-growing organoids, another 30 μL of organoid medium containing the same amounts of drug as at T=0 hours was added to the wells, and experiments were stopped at T=120 hours by assessing cell viability using Cell-Titer-Glo.

### Small Molecule Drug Screen

After identifying a predicted and in vitro confirmed resistant organoid culture to VP (TORG40), we expanded this culture to perform a high-throughput drug screen using the Ludwig Selleck Anti-Cancer Library (https://iccb.med.harvard.edu/ludwig-selleck-anti-cancer-library) containing 386 characterized bioactive compounds with validated targets against cancer-related proteins at the ICCB-Longwood Screening Facility at Harvard Medical School (HMS). Prior to screening, TORG40 was grown in 100% BME gel domes in 24-well plates supplemented with fully defined organoid medium. Two days prior to library addition, the domes of gel containing organoids were digested into small clumps using TrypLE, filtered, and resuspended in organoid medium containing 6% BME gel. Every 6 confluent wells of a 24-well plate yielded one 384-well plate. The organoid suspension was then plated into 384-well plates (Corning 3570) with 30 μL/well using the ThermoFisher Multidrop Combi Reagent Dispenser.

On the day of screening, 100 nL of each library compound in DMSO was pin-transferred into the 384-well plate. Each compound was plated in a 9-step logarithmic concentration range from 25 μM down to 0.25 nM in the well. For each drug compound at every concentration, there were four replicates. After adding the library compounds, two replicates received 10 μL/well of organoid medium, and two replicates received 10 μL/well of organoid medium containing cisplatin for a final volume of 40 μL/well and a cisplatin concentration of 5 μM. Columns 1, 2, and 23 of each plate were given 10 μL of 1 mg/ml aqueous cisplatin as a positive control, and column 24 (negative control) was given 10 μL of organoid medium with DMSO to normalize the amount of vehicle (DMSO) across the wells (< 1% DMSO/well). Column 24 in the +cisplatin plates was given organoid medium with DMSO and cisplatin to achieve a final cisplatin dose of 5 μM. Plates were stacked 4 high in a high-humidity incubator and left for three days. The library pin transfers were split into four library plates at a time due to the large number of assay plates. On day 3 after pin transfer, cells were lysed with 20 μL/well Cell Titer-Glo (1:2 ratio to medium volume, Promega, Cat. No. G7572), shaken at room temperature for 15 minutes, and viability was measured by luminescence using a Perkin Elmer EnVision.

To calculate relative cell viability, crosstalk-corrected luminescence values were normalized to the appropriate negative controls. Combination treatment wells (containing cisplatin) were normalized to negative controls containing 5 μM cisplatin to control for the moderate killing by cisplatin as a monotherapy (∼5%). Using the average relative cell count of the duplicates for each condition at different concentrations (concentrations on a logarithmic scale), dose-response curves were generated using GraphPad Prism, and the area-under-the-dose-response-curve (AUC) was calculated using base R.

The ratio of AUCs between the condition with cisplatin and without cisplatin was used to rank the compounds and select the top 5% of compounds in the screen with the largest reduction in AUC with the addition of 5 μM cisplatin (n=19). We then reviewed only those compounds of which the IC50 value of the combination treatment with cisplatin was among the smallest 20% in the compound screen (n=17/19), and chose a selection of 8 compounds, with various mechanisms of action, with a maximum of 3 compounds per drug target to validate in a secondary assay.

### Validation of top hits using an imaging-based assay

TORG40 was plated in suspension (90 μL/well) in a 96-well plate as described above. Two days after plating, drugs/controls and a fluorescent cytotoxicity dye, CellTox at a 1:1000 final concentration (Promega, Cat. No G8743), were added in 30 μL of organoid medium. Each experimental drug was tested at 3 concentrations (5 μM, 1 μM, and 0.2 μM), both with and without a fixed dose (5 μM) of cisplatin. In addition to the experimental wells, a negative control (0.8% DMSO), a negative control with 5μM cisplatin, a positive control with high dose, 100 μM cisplatin, and a positive control with lysis buffer solution (Promega, Cat. No. G8743) were included. The drugged organoids were incubated for 72 hours in the Incucyte live-cell analysis system (Sartorius, Model No. S3-C2) with images automatically taken of each well every 6 hours. The system automatically selects one focus position for imaging, which cannot be changed manually. After 72 hours, cells were lysed with Cell Titer Glo (Promega, Cat. No. G7572) at a 1:2 ratio, and luminescence readings were taken on the BioTek Synergy HT after leaving the plate on a shaker for 30 minutes. We evaluated various outcome measures acquired in Incucyte and found fluorescent object count relative to baseline (T=0) to be the most robust outcome measure. Plots were made using GRmetrics and ggplot2 in R.

### Validation of top hits in other resistant organoid cultures

TORG40, TORG58, and TORG139 were plated in suspension (90 μL/well) in a 96-well plate as described above. Similarly, two days after plating, drugs/controls and CellTox at a 1:1000 final concentration (Promega, Cat. No. G8743) were added in 30 μL of organoid medium. ABT-263 (SelleckChem, Cat. No. S1001) was tested at a single dose with and without 5 μM cisplatin. In addition to the experimental wells, positive and negative controls were included as described above. The drugged organoids were incubated for variable times ranging from 72 hours to 120 hours, depending on individual growth rates, in the Incucyte, with images automatically taken of each well every 6 hours. For cultures that were incubated for more than 72 hours, drugs and organoid medium were replenished after 72 hours (same amounts as the first time). After incubation, cells were lysed with Cell Titer Glo (Promega, Cat. No. G7572) at a 1:2 ratio, and luminescence readings were taken on the BioTek Synergy HT after leaving the plate on a shaker for 30 minutes.

### Synergy testing

A formal synergy experiment was performed with ABT-263 for TORG40, TORG58, and TORG139 by plating the organoids in suspension (90 μL/well) in a 96-well plate as described above. Similarly, two days after plating, drugs/controls and CellTox at a 1:1000 final concentration (Promega, Cat. No. G8743) were added in 30 μL of organoid medium. Cisplatin was added at 5 increasing doses (0 μM, 1 μM, 5 μM, 10 μM, 50 μM, and 100 μM) along the vertical axis and 4 increasing doses of ABT-263 (0 μM, 0.01 μM, 0.1 μM, 0.5 μM, and 1 μM) along the horizontal axis of a 96-well plate. Each dose combination was tested in technical duplicates within each plate, with two biological replicates per organoid. The drugged organoids were incubated for 72 hours in the Incucyte, with images automatically taken of each well every 6 hours. After incubation, cells were lysed with Cell Titer Glo (Promega, Cat. No. G7572) at a 1:2 ratio, and luminescence readings were taken on the BioTek Synergy HT after leaving the plate on a shaker for 30 minutes. The synergyfinder package in R was used for analyzing these multiple drug combination datasets and provided implementations for 4 popular synergy scoring models, including the Bliss synergy score(52,89). Synergy plots show the results of one biological replicate for each organoid culture.

### Quantification and Statistical Analysis

Statistical methods are outlined in the respective figure legends where applicable. Statistical analyses were performed in R. Comparisons of gene expression levels in subsets of organoids were made using Wilcoxon rank sum tests, with p-values adjusted for multiple hypothesis testing using the Benjamini-Hochberg (BH) method. P-values <0.05 were considered statistically significant. Pearson correlation analysis was used to test correlations between expression scores of selected biomarkers in organoids and clinical features of the corresponding tumors. Drug response outcomes are reported as mean ±standard error of the mean. Statistical testing for cell viability outcomes (by Cell-Titer Glo) was done using a one-way ANOVA, followed by a post-hoc Tukey test. After manual adjustments to exclude background fluorescence, setting the radius and the fluorescence threshold, the fluorescent object count normalized to baseline (T= 0 hours) values were extracted from the Incucyte integrated analysis software and subsequently visualized in R. Statistical testing for fluorescence outcomes (by CellTox) was done using a Wilcoxon test for comparing two groups, or a Kruskal-Wallis test, followed by a Dunn’s test for comparison of three or more groups.

